# Affective representations and brain–body dynamics during emotional video viewing

**DOI:** 10.64898/2026.10.28.762567

**Authors:** Weiyong Xu, Pinja Ronkainen, Tiina Parviainen

## Abstract

Emotional events evoke changing neural and bodily responses, but how these responses relate to overall emotional evaluations and how brain and body responses correspond over time remain unclear. Ninety-two participants viewed 25 silent emotional videos, each approximately one minute long, and rated valence and arousal after each video. We recorded brain activity using magnetoencephalography alongside cardiac, electrodermal, respiratory and facial-muscle activity. Representational similarity analysis (RSA) compared source-resolved cortical spectral patterns and peripheral response patterns with participants’ valence and arousal ratings, adjusting each affective dimension for the other. Across whole videos, valence similarity corresponded to beta-band power patterns predominantly in left sensorimotor and frontotemporal cortex and to facial-muscle responses. Arousal similarity corresponded to alpha-band power patterns in the right frontoparietal cortex and to phasic electrodermal activity. The neural associations persisted after adjustment for visual and semantic similarities between the videos. Windowed neural correspondence with retrospective valence was significant from around 10 s onwards, with both neural and facial-muscle associations strengthening towards the end of the videos, whereas no significant windowed arousal associations were detected throughout viewing. We further applied dynamic RSA to compare neural and physiological response patterns across videos at different temporal lags. Correspondence exceeded a temporally decoupled reference for facial muscles, heart rate and electrodermal activity, but not respiration rate. On average, corrugator activity preceded corresponding neural patterns by about 1 s, zygomaticus activity occurred at approximately the same time, and heart-rate and electrodermal activity followed about 1.7 s and 4.5 s later, respectively. Together, these findings link overall affective evaluations to complementary neural and peripheral response geometries and reveal signal-specific temporal profiles of ongoing brain–body correspondence.

## Introduction

Emotional episodes unfold across brain and body, involving changes in neural activity, autonomic physiology, facial expression and subjective feeling. Componential accounts describe emotion as an interaction among appraisal, action preparation, bodily regulation, expression and feeling (Sander et al., 2018), with coordination that varies across people, emotions and contexts (Mauss et al., 2024). Empirical findings likewise show partial coherence: subjective and autonomic responses share some structure, while much autonomic variation does not covary with subjective experience (Cuve et al., 2023). A central challenge is therefore to understand whether episodes that are experienced as similar also evoke similar neural and bodily responses, and how these relationships develop over time.

Naturalistic videos allow these relationships to be examined during extended emotional episodes. Their evolving perceptual information, actions and event structure engage brain systems that integrate information over multiple timescales (Hasson et al., 2008; Jääskeläinen et al., 2021; Saarimäki, 2021). The resulting experiences can be characterized in terms of valence, describing pleasantness–unpleasantness, and arousal, describing activation–deactivation (Russell, 1980). These two dimensions emerge consistently in emotional experiences elicited by videos, narratives and everyday events (Han et al., 2024). Because such experiences vary along continuous gradients rather than forming separate clusters (Cowen & Keltner, 2017; Han et al., 2024), valence and arousal offer a compact and general basis for comparing emotional episodes. Extended episodes also require distinguishing responses during an event from its subsequent overall evaluation. Studies combining continuous and retrospective reports show that overall judgments need not weight every moment equally, with the ending of an emotional sequence sometimes receiving disproportionate weight (Fredrickson & Kahneman, 1993; Li & Lapate, 2024). This distinction motivates asking when neural and bodily response patterns during viewing correspond to the evaluation subsequently assigned to the episode.

Functional magnetic resonance imaging (fMRI) studies using multivariate methods show that affective experiences are reflected in distributed neural patterns. Valence-related patterns in orbitofrontal cortex generalize across static pictures and tastes (Chikazoe et al., 2014), while emotion categories elicited by short films or mental imagery can be distinguished from activity in prefrontal, sensorimotor and posterior midline regions (Saarimäki et al., 2016). Other film-based fMRI studies identify high-dimensional affective representations across association cortex and modality-independent representations in prefrontal and temporal regions (Horikawa et al., 2020; Lettieri et al., 2024). Complementing this spatial evidence, magnetoencephalography (MEG) reveals the rapid emergence of affect-related information: neural similarity during picture viewing corresponds to independently obtained valence and arousal ratings within the first 200 ms, with associations peaking around 300 ms (Grootswagers et al., 2020). Electroencephalography (EEG) further links high-arousal pictures to reduced alpha- and lower-beta power (Schubring & Schupp, 2021), and momentary arousal during movies to reduced low-frequency power in occipitoparietal regions (Camenzind et al., 2026). Together, these studies establish spatial, temporal and spectral aspects of affective processing. Less clear is how participant-specific evaluations relate to cortical spectral patterns throughout extended emotional episodes, and how this organization compares with concurrent bodily responses.

Bodily responses also reflect aspects of emotional experience, although their relationship with subjective feeling differs across physiological systems. Facial-muscle activity is associated with valence (Larsen et al., 2003), whereas electrodermal responses relate to arousal (Kreibig, 2010; Lang et al., 1993). These regularities coexist with substantial physiological variation within and across emotional states (Siegel et al., 2018). During emotional films, correspondence with reported feelings differs between facial and autonomic measures and changes across phases of the episode (Constantinou et al., 2023). Cardiac and electrodermal responses also show temporal alignment across viewers of emotional films, with the consistency of these responses varying along the film’s emotional timeline (Golland et al., 2014). Representational analyses further show that affective–physiological correspondence is selective across signals and induction contexts (Ventura- Bort et al., 2022), and that its characterization depends on how within-person response variability is represented (Koppold et al., 2026). The relevant question is therefore not whether bodily activity provides a single signature of emotion, but which physiological responses preserve the distinctions between episodes expressed in a participant’s evaluations, and whether that correspondence is sustained during viewing.

Representational similarity analysis (RSA) provides a common framework for relating subjective ratings to neural and bodily responses. RSA characterizes each domain through pairwise dissimilarities among the same events, allowing cortical activity patterns, individual physiological signals and ratings to be compared despite their different units and dimensionalities (Diedrichsen & Kriegeskorte, 2017; Kriegeskorte et al., 2008). Applied to emotional videos, it asks whether episodes evaluated similarly also evoke similar neural or bodily responses. Furthermore, RSA can incorporate stimulus properties that contribute to both neural activity and emotional evaluation. For example, representations from object-recognition models and object-selective visual cortex predict emotion ratings across diverse everyday scenes (Gao et al., 2025). Controlling for measured video features, such as luminance, motion and visual or semantic content, can therefore help assess whether neural correspondence with subjective evaluations persists beyond these stimulus properties.

Beyond their relationship with overall affective evaluations, neural and bodily responses may also correspond as an emotional episode unfolds. Recurrent pathways link neural activity with bodily states through central regulation and ascending visceral and somatosensory signals (Azzalini et al., 2019; Barrett & Simmons, 2015; Critchley & Harrison, 2013). Simultaneous neural and cardiac recordings associate emotional arousal with changing brain–heart interactions and cortical responses to heartbeats (Candia-Rivera et al., 2022; Fourcade et al., 2024). Electrodermal activity has been related to orbitofrontal and insular activity using fMRI (Critchley et al., 2000) and to early occipitotemporal MEG responses during emotional picture viewing (D’Hondt et al., 2010). Facial EMG covaries with inferior frontal and supplementary motor activity (Likowski et al., 2012), while experimentally reducing frown-muscle feedback attenuates amygdala responses during imitation of angry expressions (Hennenlotter et al., 2009). During natural breathing, MEG and intracranial EEG also reveal respiratory-phase modulation of oscillatory activity across cortical and limbic regions (Kluger & Gross, 2021; Zelano et al., 2016). These relationships span different timescales: facial reactions can emerge within hundreds of milliseconds, whereas electrodermal responses develop over several seconds (Bach et al., 2010; Dimberg & Thunberg, 1998). Examining only simultaneous activity may therefore overlook correspondence at different temporal lags. To examine these temporal relationships, we adopted the dynamic RSA approach initially developed by de Vries and Wurm (2023). During dance-video viewing, they compared MEG response patterns with time-varying representations of visual features, body posture and motion, distinguishing neural responses that followed the input from representations that anticipated upcoming movements. Here, we adapted this approach to compare neural and physiological response patterns across emotional videos at different temporal lags, assessing correspondence with bodily responses occurring earlier, simultaneously or later.

In this study, we recorded MEG alongside cardiac, electrodermal, respiratory and facial-muscle activity in 92 participants viewing 25 silent emotional videos. After each clip, participants rated their experienced valence and arousal. Using RSA, we tested whether neural and bodily responses share aspects of affective organization while differing in their temporal expression. Whole-video analyses assessed whether similarities in valence and arousal were reflected in cortical spectral patterns and peripheral responses, with particular interest in facial-muscle correspondence with valence and electrodermal correspondence with arousal. Windowed analyses examined whether correspondence with retrospective evaluations was sustained during viewing and how it evolved as each episode unfolded. Finally, dynamic RSA tested whether ongoing brain–body correspondence depended on temporal alignment and exhibited different spatiotemporal profiles across physiological systems. Together, these analyses examine how neural and bodily responses relate to subjective evaluations and to one another as emotional episodes unfold.

## Methods

### Participants

Ninety-two participants (66 female; age range 18–56 years, *M* = 29.3, *SD* = 8.9) were recruited through mailing lists, online advertisements, and posters. Inclusion criteria were age between 18 and 60 years; normal or corrected-to-normal vision; absence of any diagnosed severe psychiatric disorder (e.g., major depressive disorder); no neurological illness or use of medication affecting the central nervous system; and no metal implants incompatible with MEG recording. Individuals with severe psychiatric conditions were excluded to minimize potential risk to those with pre-existing psychological vulnerability. Participants received a €15 gift card or their choice of chocolate or coffee as compensation. The study was approved by the Ethics Committee of the University of Jyväskylä and conducted in accordance with the Declaration of Helsinki. All participants provided written informed consent before participation.

### Stimuli and task

Participants viewed 25 silent video clips (approximately 60 s each; 1280 × 720 pixels), comprising five clips from each of five intended emotion categories: anxiety/fear, sadness, neutral, contentment, and happiness. Seven clips were selected from the FilmStim database (Schaefer et al., 2010). The remaining clips were selected by the research team and evaluated in two rounds of validation surveys. The selection criteria are described in the Supplementary methods (“Stimulus selection criteria”). The video task formed part of a larger experimental session that also included resting-state recordings, an escape-game task (Pultsina et al., 2026), and a modified Trier Social Stress Test (Figure 1A).

**Figure 1.**
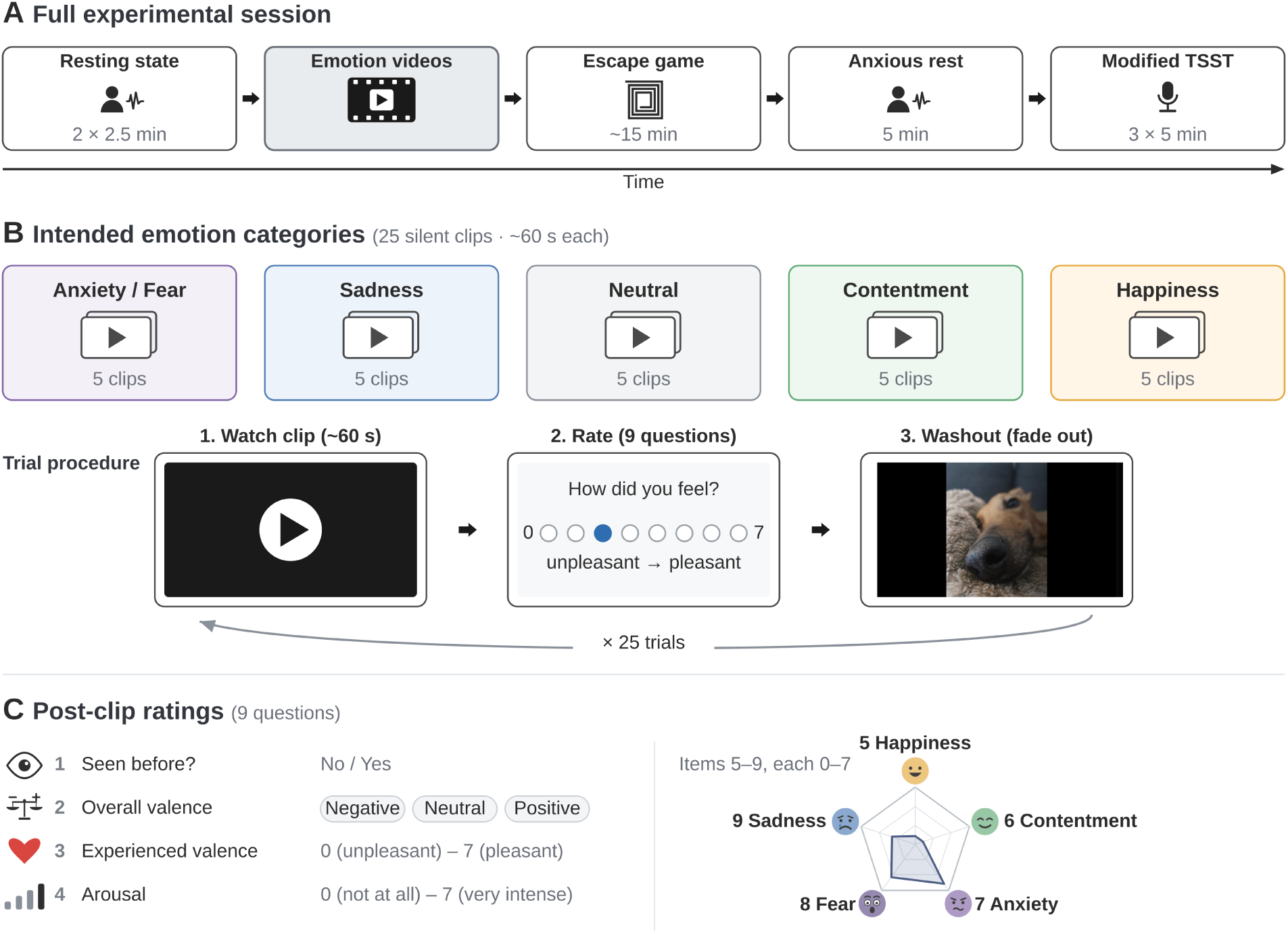
Experimental design. **(A)** Overview of the recording session. Participants completed resting-state recordings, the emotion video-viewing task used in the current study (highlighted), an escape-game paradigm, an anxious resting-state block, and a modified Trier Social Stress Test (TSST). **(B)** Top: the stimulus set comprised 25 silent video clips (∼60 s each; 1280 × 720 px), with five clips from each of five intended emotion categories. Bottom: on each trial participants passively viewed one clip, answered nine self-paced questions, and then saw a washout image that gradually faded to black before the next trial; no fixation cross was used. Category icons and the video-viewing and rating screens are schematic; the washout screen shows the original washout image. **(C)** The nine post-clip ratings and their response formats: prior familiarity, overall valence category, experienced valence, arousal, and the strength of five discrete emotions (0–7 each), which together form an emotion profile per clip (radar plot). Highlighted responses illustrate one hypothetical negative clip and are schematic, not empirical data. Questions were presented in Finnish (see the Supplementary methods).

On each trial, participants passively viewed one video clip and then answered nine self-paced questions presented in Finnish (see the Supplementary methods). These assessed prior familiarity, overall valence category (positive, negative, or neutral), experienced valence (0 = very unpleasant, 7 = very pleasant), perceived video intensity (0 = not at all intense, 7 = very intense), and five discrete emotions: happiness, contentment, anxiety, fear, and sadness (0 = not at all, 7 = very much). Valence, intensity, and discrete emotions were rated on eight-point scales. Perceived intensity served as the arousal measure throughout this study. Between trials, a washout image gradually faded to black; no fixation cross was used. A break of approximately 1 min was provided between trials 12 and 13. All participants viewed the clips in the same fixed order.

### Data acquisition

MEG was recorded using a 306-channel Elekta Neuromag TRIUX system (102 magnetometers and 204 planar gradiometers) in a magnetically shielded room at the Centre for Interdisciplinary Brain Research (CIBR), University of Jyväskylä. Participants were seated with the gantry positioned at 68°. Signals were sampled at 1000 Hz with an online band-pass filter of 0.1–330 Hz. Head position was monitored continuously using five head-position indicator coils, with three on the forehead and one behind each ear. Before recording, the nasion, left and right preauricular points, and coil locations were digitized using a Polhemus Isotrak system, with a target of 200 additional head-shape points.

The electrooculogram (EOG) was recorded using two electrodes positioned diagonally around the eyes. Peripheral recordings comprised the electrocardiogram (ECG), electrodermal activity (EDA), facial electromyography (EMG), and respiration. Bipolar EMG recordings targeted the left corrugator supercilii and zygomaticus major muscles. ECG electrodes were positioned diagonally across the chest, approximately 1–2 cm below the right clavicle and 9–10 cm below the left clavicle. EDA was recorded using Ambu NEUROLINE 710 electrodes placed on the medial plantar surface of the right foot. Respiration was recorded primarily using a custom-built respiratory belt (Karjalainen et al., 2025); a commercial belt (Spes Medica) was used for 26 participants because of technical problems with the custom-built belt. Both belt types were positioned over clothing around the lower chest.

Structural MRI was acquired for 91 of the 92 participants using a 1.5 T GE Signa HDxt scanner at SYNLAB Jyväskylä. T1-weighted images were collected with a three-dimensional FSPGR BRAVO sequence (TI = 450 ms, flip angle = 12°, slice thickness = 1.2 mm, slice spacing = 0.6 mm). For the participant without an individual MRI, the FreeSurfer fsaverage template (Fischl, 2012) was uniformly scaled to the digitized head shape.

### MEG preprocessing

MEG preprocessing was performed using MNE-Python (version 1.11.0; Gramfort et al., 2013). Noisy and flat channels were first identified using the automated Maxwell-filter-based bad-channel detection procedure implemented in MNE-Python (find_bad_channels_maxwell). The resulting channel selections were then reviewed manually. For each participant, bad-channel lists were combined across the video task, eyes-open resting-state, and empty-room recordings so that the same channels were marked as bad when processing these recordings.

Temporal signal-space separation (tSSS; Taulu & Simola, 2006), incorporating fine-calibration and cross-talk corrections, was applied to suppress external magnetic interference. Head movements were compensated using the continuously recorded head-position estimates, with the mean head position during the video task serving as the reference position. The tSSS procedure used a 10-s temporal window and a correlation threshold of 0.98. This threshold was reduced to 0.90 for nine recordings identified as particularly noisy. The corresponding empty-room recordings were processed with matched tSSS settings to provide noise estimates for subsequent source reconstruction.

Following tSSS, the recordings were visually inspected, and noisy segments were marked for exclusion. Cardiac and ocular artifacts were identified using independent component analysis (ICA) with the FastICA algorithm. ICA was fitted to a copy of the video-task recording filtered at 1–40 Hz, with the number of components determined by a 99% explained-variance threshold. Automated EOG- and ECG-based component scores guided visual inspection of component topographies and time courses, and components judged to reflect artifacts were selected manually for removal. The ICA solution was then applied to the continuous recordings filtered at 0.1–100 Hz using a zero-phase finite impulse response filter. Video onset times were shifted forward by 23 ms to account for the measured trigger-to-display delay.

### Source reconstruction

Individual cortical surfaces were reconstructed using FreeSurfer 8.1.0 (Dale et al., 1999), with T1-weighted images and, where available, T2-weighted images for pial-surface refinement. For participants with individual MRI, automated coregistration aligned the digitized anatomical landmarks and then refined the fit to the head-shape points using iterative closest-point fitting, excluding points more than 5 mm from the scalp between fits (Houck & Claus, 2020). A single-layer boundary-element model of the inner skull, with conductivity set to 0.3 S/m, was used. Candidate sources were distributed over the cortical white-matter surface at ico-4 spacing. Cortical activity was estimated using dynamic statistical parametric mapping (dSPM; Dale et al., 2000). Noise covariance was estimated from each participant’s preprocessed empty-room recording using shrinkage estimation. The inverse solution used loose orientation constraints of 0.2, depth weighting of 0.8, and regularization λ² = 1.0. Source time courses retained the signed component normal to the cortical surface. Covariance estimation and inverse modeling accounted for the data rank following Maxwell filtering.

### Spectral estimation

Spectral power was estimated after resampling the MEG recordings to 250 Hz. For both whole-video and windowed spectral analyses, power was estimated in native source space and mapped to the fsaverage ico-4 template before logarithmic transformation. For the whole-video analysis, power spectral density was estimated over the first 59 s of each video using 1.6-s Hann windows with 50% overlap. Log-transformed power was averaged within five frequency bands: theta (4–8 Hz), alpha (8–13 Hz), beta (13–30 Hz), low gamma (30–45 Hz), and high gamma (60–90 Hz). To examine changes during viewing, spectra were estimated using a single Hamming-tapered periodogram within each of 29 consecutive, non-overlapping 2-s windows spanning the first 58 s of each video. Windowed analyses used log-transformed power at 37 frequencies spanning 4–90 Hz on a logarithmic scale, excluding 45–60 Hz, rather than averages within frequency bands.

Spectral analyses excluded intervals marked as bad during visual inspection and muscle-artifact intervals lasting at least 100 ms. Muscle artifacts were detected using MNE-Python’s annotate_muscle_zscore procedure, applied to magnetometer signals in the 55–95 Hz band with a muscle-score threshold of 5. For whole-video power spectra, spectral windows overlapping excluded intervals were discarded, and estimates were computed from the remaining windows within each clip; clips with no usable windows were omitted. For time-resolved spectral analyses, any 2-s window overlapping an excluded interval was rejected.

### Peripheral physiological signals

ECG, EDA, and respiration recordings were processed using NeuroKit2 (Makowski et al., 2021). Corrugator supercilii and zygomaticus major EMG signals were notch-filtered at 50 Hz and band-pass filtered between 20 and 300 Hz using a fourth-order Butterworth filter. Signals were then full-wave rectified and low-pass filtered at 8 Hz using a second-order Butterworth filter. Filters were applied in forward and reverse directions to avoid phase shifts. Mean envelope amplitude was calculated for each video. ECG processing comprised signal cleaning, R-peak detection, and derivation of a continuous heart-rate time course from successive interbeat intervals. Each video was summarized by mean heart rate and heart-rate variability (HRV), quantified as the root mean square of successive differences between interbeat intervals (RMSSD). Heart rate was retained for time-resolved analyses, whereas RMSSD was calculated only at the whole-video level. The continuous EDA signal was low-pass filtered at 3 Hz using a fourth-order Butterworth filter, and a phasic signal was obtained using a second-order Butterworth high-pass filter at 0.05 Hz. Both digital filters were applied in forward and reverse directions to avoid phase shifts. Because the recording system applied a 0.1-Hz analog high-pass filter, this signal represented relative electrodermal fluctuations in arbitrary units rather than absolute skin-conductance levels. Respiration processing identified respiratory peaks and troughs and yielded a continuous respiration-rate time course, from which mean respiration rate was calculated for each video.

Physiological recordings underwent visual inspection and signal-specific quality checks to identify recording artifacts and unreliable cardiac or respiratory cycle detection. Exclusions were applied separately by modality and facial-muscle channel. The retained samples comprised 92 participants for corrugator EMG, 91 for zygomaticus EMG, 79 for ECG, 80 for EDA, and 86 for respiration. Continuous physiological signals were resampled to 50 Hz for the dynamic brain–body analysis.

### Characterization of subjective ratings

Subjective ratings were characterized to assess the emotional range of the stimulus set and its correspondence with the intended emotion categories. Mean ratings of the five discrete emotions were summarized for each category and individual video to describe their emotional profiles and overlap (Figure 2A). The proportions of negative, neutral and positive valence judgments were calculated within each category to assess whether its videos were experienced as unpleasant, neutral or pleasant (Figure 2B). To compare valence and arousal across categories, ratings were averaged across the five videos in each category within each participant (Figure 2C, top and middle). The relationship between clip-mean valence and arousal was examined to assess how the two dimensions varied together across the stimulus set (Figure 2C, bottom). Finally, agreement in valence, arousal and discrete-emotion ratings across participants was assessed to establish how consistently individual videos were evaluated (Figure 2D). Subsequent RSA used each participant’s ratings for individual videos.

**Figure 2.**
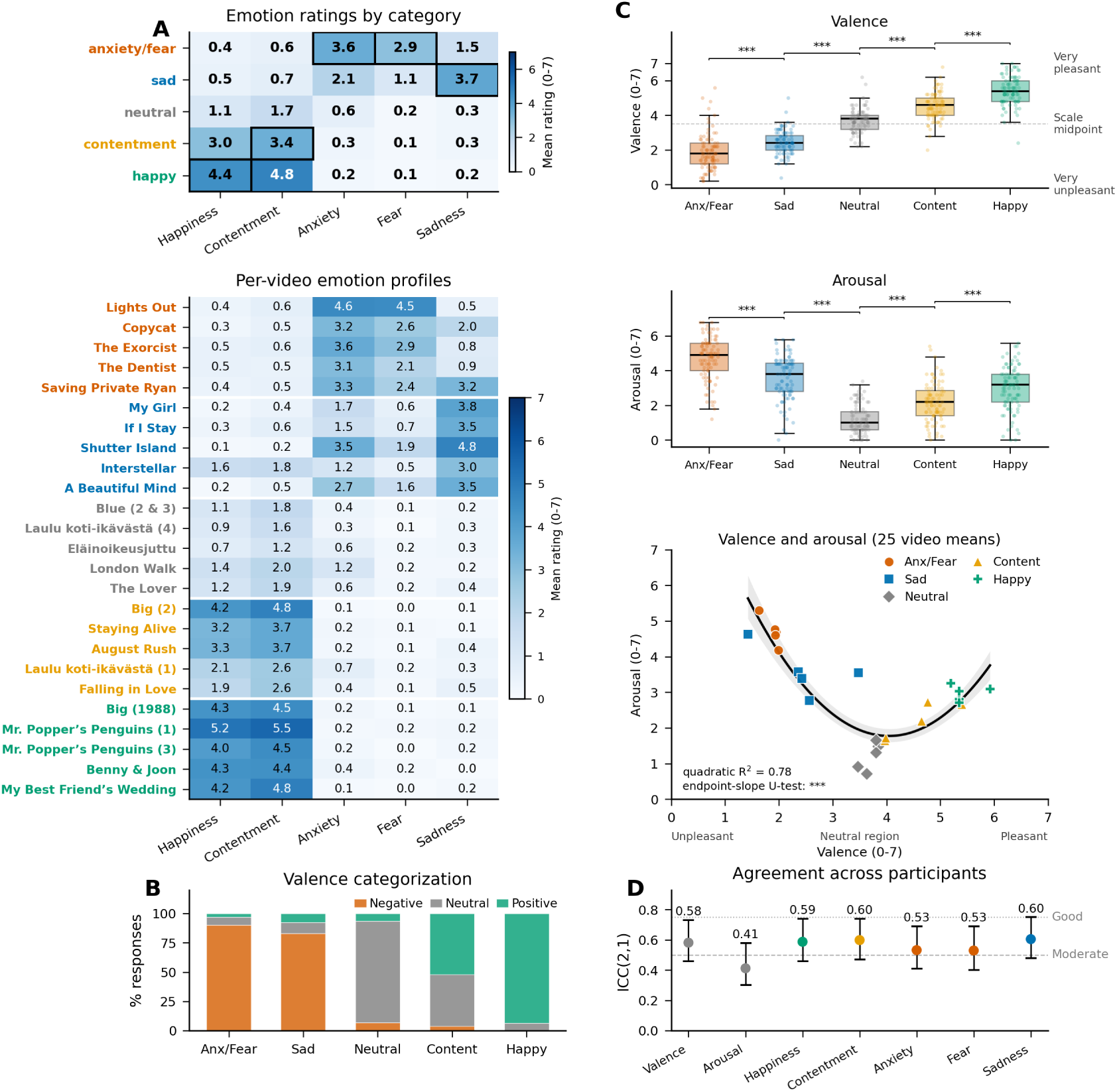
Subjective emotional ratings of the videos. Ratings from 92 participants viewing 25 videos, with five videos per emotion category. **(A)** Mean discrete-emotion ratings by category (top) and individual video (bottom). Cell values and colour indicate mean ratings on a 0–7 scale. Black outlines mark the intended dominant emotions; white dividers separate video categories. **(B)** Percentages of responses categorized as negative, neutral, or positive within each video category. **(C)** Valence and arousal ratings by category (top and middle). Dots represent individual participants’ means across five videos. Valence ratings ranged from 0 (very unpleasant) to 7 (very pleasant); the dashed line marks the scale midpoint (3.5). Boxes span the interquartile range, horizontal lines mark medians, and whiskers extend to the most extreme observations within 1.5 interquartile ranges of the box edges. Brackets indicate adjacent-category comparisons using two-sided paired Wilcoxon signed-rank tests, with Benjamini–Hochberg FDR correction across all eight comparisons (***, *q* < .001). Bottom: mean valence and arousal for each video, with category indicated by colour and marker shape. The curve shows a quadratic fit (*R*² = .78); shading indicates a pointwise 95% confidence band from 5,000 participant bootstrap samples (endpoint-slope U-test: ***, *p* < .001). **(D)** Agreement across participants, quantified using the two-way random-effects, absolute-agreement, single-measure intraclass correlation coefficient, ICC(2,1). Points and error bars show estimates and 95% confidence intervals. Dashed and dotted lines mark conventional reference thresholds for moderate (.50) and good (.75) agreement (Koo & Li, 2016).

### RDM construction

#### Stimulus-feature RDMs

To account for visual properties that potentially covary with affective ratings, we constructed four stimulus-feature representational dissimilarity matrices (RDMs), describing mean luminance, motion-energy patterns, early visual-network features, and high-level semantic features. Features were represented at 5 Hz. For whole-video analyses, features were averaged over the first 59 s of each video, matching the MEG analysis interval. For windowed analyses, features were averaged separately within each of the 29 consecutive, non-overlapping 2-s windows spanning the first 58 s. Pairwise distances between videos’ feature vectors yielded one 25 × 25 RDM per model for the whole-video analysis and one per model and window for the windowed analysis. Mean luminance was calculated as the average grayscale pixel intensity. Motion-energy features were extracted using a bank of spatiotemporal Gabor filters implemented in pymoten, capturing patterns across spatial locations, motion directions, and spatial and temporal frequencies (Nunez-Elizalde et al., 2021). Videos were resampled to 24 frames per second for motion-energy extraction, and filter responses were aggregated into 5-Hz bins. Early visual features were obtained from the first residual block of an ImageNet-pretrained ResNet-18 (He et al., 2016), with spatial averaging yielding 64 features per frame. High-level semantic features were represented by 512-dimensional image embeddings from an OpenCLIP ViT-B/16 model pretrained on LAION-2B (Radford et al., 2021; Schuhmann et al., 2022). Luminance dissimilarities were calculated as absolute differences after standardization across videos. Motion-energy, early visual, and semantic dissimilarities were calculated using correlation distance (1 − Pearson’s *r*) between video-level feature vectors.

#### Subjective-affect RDMs

For each participant, subjective-affect RDMs were constructed from the post-video ratings. Valence and arousal dissimilarities were defined as the absolute difference between each pair of videos’ experienced-valence and perceived-intensity ratings, respectively, yielding a separate 25 × 25 RDM for each dimension. Primary analyses focused on valence and arousal. Three additional RDMs were examined descriptively but not included in the partial-RSA models: emotion profile, calculated from Euclidean distances between the five discrete-emotion ratings; temporal proximity, calculated from absolute differences between video-onset times; and familiarity, calculated from absolute differences between prior-familiarity responses. Candidate stimulus and rating RDMs are shown in Supplementary Figure S1.

#### Physiological RDMs

For each participant, separate physiological RDMs were constructed for mean heart rate, heart-rate variability (RMSSD), mean positive phasic EDA, respiration rate, and corrugator and zygomaticus EMG. For both whole-video and windowed body–rating analyses, mean positive phasic EDA was defined as the average of the phasic signal after setting negative values to zero, with zeroed samples retained in the average. Each physiological feature was standardized across available videos within participant, and dissimilarities were calculated as absolute differences between video-level values. For windowed analyses, mean heart rate, mean positive phasic EDA, respiration rate, and facial EMG were calculated within consecutive 2-s windows, and absolute differences between videos yielded one RDM per signal and window. RMSSD was included only in the whole-video analysis. Relationships among physiological RDMs were examined descriptively (Supplementary Figure S3).

### Representational similarity analysis

Associations of neural and physiological response patterns with affective ratings were quantified by comparing the upper triangles of response RDMs with participant-specific valence and arousal RDMs. Comparisons used matching video pairs with valid response, rating, and control data. Brain–rating RSA required at least 20 usable videos for each whole-video or window-specific estimate. The primary analyses used partial Spearman correlations to assess valence while controlling for arousal, and arousal while controlling for valence. Partial correlations were calculated by removing the contribution of the control RDM from both rank-transformed RDM vectors and correlating the residuals. Unadjusted body–rating associations were examined as sensitivity analyses (Supplementary Figure S4). For the brain analyses, an additional stimulus-adjusted analysis controlled for mean luminance, motion-energy patterns, early visual-network features, and high-level semantic features (Supplementary Figure S2).

#### Whole-video RSA

Whole-video brain–rating RSA assessed local spatial patterns of source spectral power within cortical searchlights with a 20-mm geodesic radius, separately for each frequency band. Each video was represented by its spatial pattern of band-averaged log power across vertices within a searchlight. Pairwise correlation distances (1 − Pearson’s *r*) between these patterns yielded a neural RDM for each participant, searchlight, and frequency band. Neural RDMs were compared with participant-specific rating RDMs using the partial Spearman correlations described above. Whole-video body–rating RSA evaluated six physiological indices separately: heart rate, heart-rate variability, mean positive phasic EDA, respiration rate, and corrugator and zygomaticus EMG. Whole-video physiological features were calculated over the interval from video onset to the onset of the post-video questions.

#### Time-resolved RSA

Windowed brain–rating and body–rating RSA used 29 consecutive, non-overlapping 2-s windows spanning the first 58 s of each video. Because ratings were collected once after each video, participant-specific rating RDMs remained fixed across windows. The analysis therefore assessed when neural or physiological response patterns corresponded to the subsequent overall evaluation. To improve the signal-to-noise ratio of estimates from short windows, windowed brain–rating RSA combined spatial and spectral information: each video was represented by a single feature vector containing log-power values across all vertices within a Desikan–Killiany cortical parcel (Desikan et al., 2006) and the 37 retained frequency bins. These feature vectors were constructed separately for each participant, parcel, and 2-s window. Pairwise correlation distances (1 − Pearson’s *r*) between videos’ feature vectors yielded one neural RDM per participant, parcel, and window. Comparing these neural RDMs with participant-specific rating RDMs using partial Spearman correlations yielded the corresponding association estimates. For body–rating RSA, window-specific associations were calculated for heart rate, mean positive phasic EDA, respiration rate, and the two facial-muscle signals. Heart-rate variability was evaluated only at the whole-video level.

#### Dynamic brain–body RSA

Dynamic brain–body RSA examined temporal correspondence between neural and physiological response patterns without reference to affective ratings. Sign-aligned mean source time courses were extracted from each participant’s 68 Desikan–Killiany cortical parcels, with corresponding parcels matched across participants. These time courses were compared with heart rate, phasic EDA, respiration rate, and corrugator and zygomaticus EMG. Signals were sampled at 50 Hz and standardized within each video, so the analysis assessed within-video fluctuations rather than differences in overall response magnitude between videos. EDA retained its signed phasic waveform, and facial EMG was not linearly detrended.

Following the temporal sampling approach of dynamic RSA (de Vries & Wurm, 2023), a random anchor time was drawn independently within each of the 25 videos. Neural activity was averaged over a 0.2-s window centred on each anchor, and physiological activity over a 1-s window centred at a specified lag relative to that anchor. Pairwise absolute differences across videos formed a neural RDM for each parcel and a physiological RDM for each signal; these RDMs were compared using Spearman correlation. Estimates were averaged over 2,000 repetitions at each lag, from −12 to +12 s in 0.5-s steps. Lag was defined as physiological time minus neural time, so positive lags paired neural activity with later physiological activity. To isolate correspondence associated with temporal alignment, each neural RDM was also compared with a physiological RDM constructed from independently sampled anchors within the same videos. The observed correlation minus this temporally decoupled reference was the measure entered into group analysis.

### Statistical analysis

#### Subjective ratings

Adjacent categories were compared using two-sided paired Wilcoxon signed-rank tests, with Benjamini–Hochberg false-discovery-rate correction across the eight comparisons (four per rating dimension). These adjusted *p* values are denoted by *q* throughout. The relationship between clip-mean valence and arousal was described using quadratic ordinary least squares regression and assessed with an endpoint-slope U-test (Lind & Mehlum, 2010); confidence bands were obtained from 5,000 participant bootstrap samples. Agreement across participants was quantified using a two-way random-effects, absolute-agreement, single-measure intraclass correlation coefficient, ICC(2,1), with 95% confidence intervals (Koo & Li, 2016).

#### Brain–rating RSA

For both whole-video and windowed brain–rating RSA, participant correlations were Fisher-*z* transformed and tested against zero using two-sided sign-flip permutation tests with threshold-free cluster enhancement (TFCE; Smith & Nichols, 2009). Each of 2,048 permutations applied one sign to each participant’s entire map. The maximum TFCE statistic controlled family-wise error across cortical vertices within each whole-video model-by-frequency-band map, and jointly across cortical parcels and adjacent windows within each windowed model. TFCE used height and extent exponents of 2 and 0.5, respectively, with a threshold increment of 0.2; corrected *p* < .05 defined significance. Tests were conducted separately for each model and, in the whole-video analysis, each frequency band. For windowed analyses, group *t* statistics used all participants with available data at each parcel and window. Missing windows remained missing during sign-flipping, so a participant could contribute to other windows even when some were unavailable.

#### Whole-video body–rating RSA

Group inference used 10,000 shared video-label permutations. In each permutation, the same reordering of the 25 videos was applied to every participant’s physiological RDM, while rating RDMs remained fixed, and the associations were recalculated. Observed group-mean correlations were evaluated using two-sided permutation tests, with Benjamini–Hochberg false discovery rate correction across the twelve physiological-index-by-rating comparisons.

#### Windowed body–rating RSA

Group inference used the TFCE settings described above, applied along time within each signal to *t* statistics from Fisher-*z*-transformed correlations. The null distribution used 10,000 shared video-label permutations, applying the same reordering across participants, physiological signals, and windows. The maximum TFCE statistic controlled family-wise error across all windows and five signals within each affective dimension (two-sided corrected *p* < .05).

#### Dynamic brain–body RSA

Group inference used a two-sided parcel-by-lag TFCE sign-flip permutation test, with cortical parcel adjacency and 2,048 permutations. The maximum TFCE statistic controlled family-wise error across parcels and lags within each physiological signal, followed by Bonferroni correction across the five signals. Relative timing was summarized by the centroid along the lag axis, weighting significant cells by the absolute unenhanced *t* statistic.

Confidence intervals were calculated using 10,000 participant bootstrap resamples while holding the detected significant extent fixed.

## Results

### Subjective emotional ratings of the videos

Subjective ratings broadly reflected the intended video categories, with overlapping happiness and contentment responses to positive videos (Figure 2A). Categorical valence judgments were predominantly negative for anxiety/fear and sad videos, neutral for neutral videos, and positive for happy videos; contentment videos elicited a mixture of positive and neutral judgments (Figure 2B). Mean valence increased from anxiety/fear through sad, neutral, and contentment to happy videos, whereas arousal was highest for anxiety/fear videos and lowest for neutral videos (all adjacent-category comparisons for both dimensions, *q* < .001; Figure 2C). Across the 25 videos, mean arousal showed a U-shaped relationship with mean valence (quadratic *R*² = .78; endpoint-slope test, *t*(22) = 5.56, *p* < .001). Agreement across participants was lower for arousal (ICC = .41) than for valence and discrete-emotion ratings (ICCs = .53–.60; Figure 2D). The stimulus-level relationship between valence and arousal motivated the partial-RSA analyses used to assess their distinct contributions.

### Relationships among the representational models

The valence and arousal RDMs showed the strongest correspondence among the six models (Spearman ρ = .61 for the group-average RDMs; Figure 3A). Within participants, valence–arousal RDM correlations were lower (median ρ = .23, IQR = .10–.35). Correlations between the group-average affective RDMs and the four stimulus-feature RDMs were weaker (ρ = −.02–.23), indicating that the affective and visual models captured partly distinct patterns of differences between videos. Multidimensional scaling provided a descriptive visualization of these relationships (Figure 3B). At the participant level, variance inflation factors calculated from ranked RDM entries were low: model-wise medians ranged from 1.10 to 1.41, with a maximum of 1.57 across all participants and models (Figure 3C). These diagnostics indicated limited multicollinearity in the six-model design used for the stimulus-adjusted analyses. The emotion-profile RDM showed substantial overlap with valence across participants (median Spearman ρ = .70, IQR = .57–.78; Supplementary Figure S1D). This overlap supported retaining emotion profile as a supplementary descriptive model while focusing the main analyses on valence and arousal.

**Figure 3.**
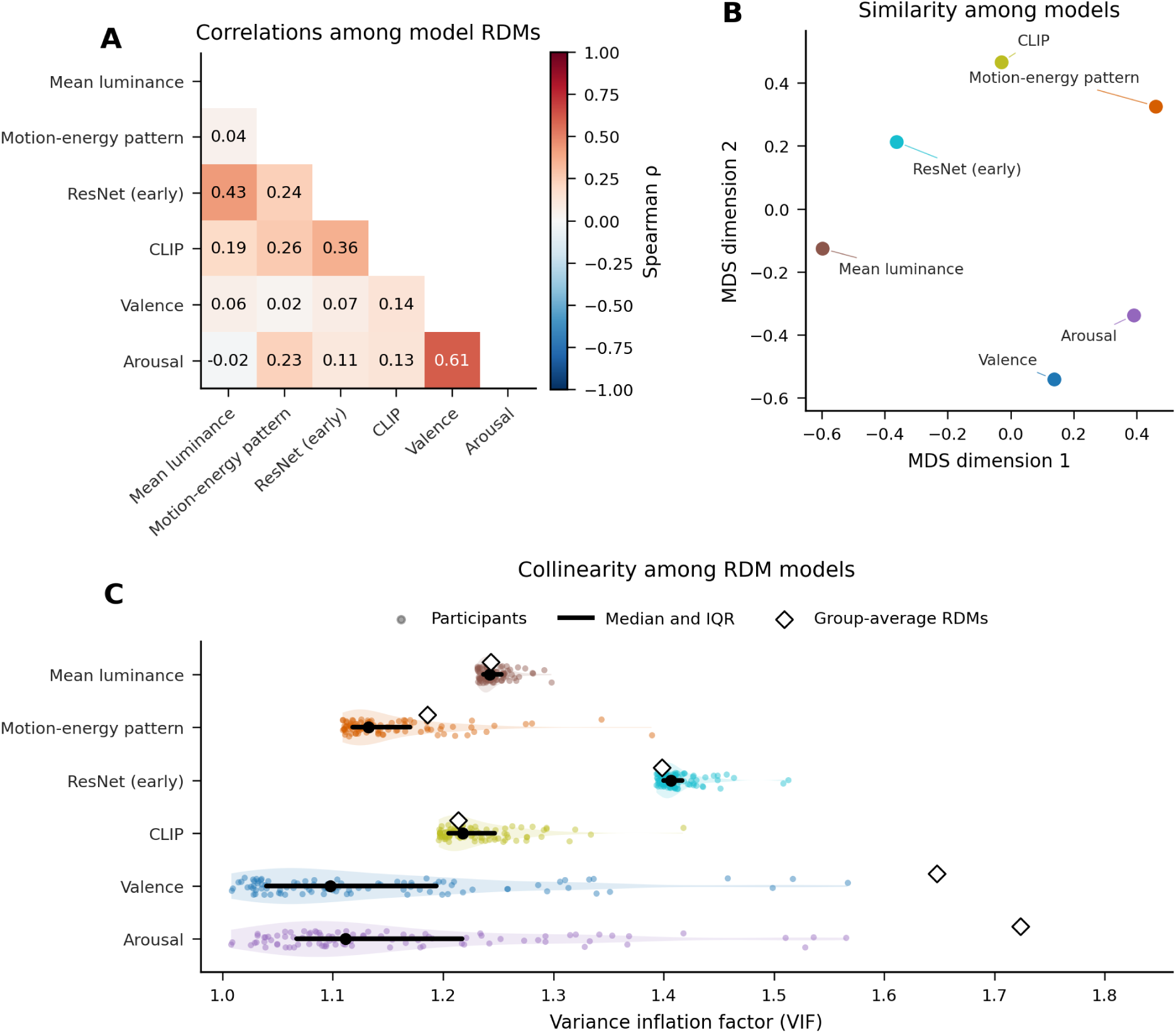
Relationships among the representational models. **(A)** Spearman correlations among the six RDMs used in the stimulus-adjusted RSA. Valence and arousal RDMs were averaged across 92 participants before calculating correlations; stimulus-feature RDMs were shared across participants. Correlations were calculated across the 300 unique video pairs, with self-correlations omitted. ResNet (early) denotes early visual-network features, and CLIP denotes high-level semantic features. **(B)** Two-dimensional metric multidimensional scaling (MDS) of model dissimilarities, defined as 1 − ρ from panel A. Distances approximate model dissimilarities; axis orientation and the origin are arbitrary (normalized Stress-1 = 0.148). **(C)** Variance inflation factors (VIFs) calculated from ranked RDM entries separately for each participant. Each VIF quantifies one model’s collinearity with the other five. Coloured dots represent participants (*n* = 92), pale shading shows their distributions, and black points and bars indicate medians and interquartile ranges. Open diamonds show VIFs calculated from the group-average RDMs rather than averages of participant-level VIFs.

### Neural representations of valence and arousal

Whole-video neural dissimilarities were associated with valence in the beta band and arousal in the alpha band after adjusting each affective dimension for the other (Figure 4A). The valence– beta association survived spatial TFCE correction, predominantly in left precentral, postcentral, inferior frontal, and temporal cortex. The arousal–alpha association was mainly in right frontal and parietal cortex, particularly the precentral, postcentral, and supramarginal regions (minimum corrected *p* < .001 for both associations). The arousal–beta association did not survive correction (minimum corrected *p* = .054).

**Figure 4.**
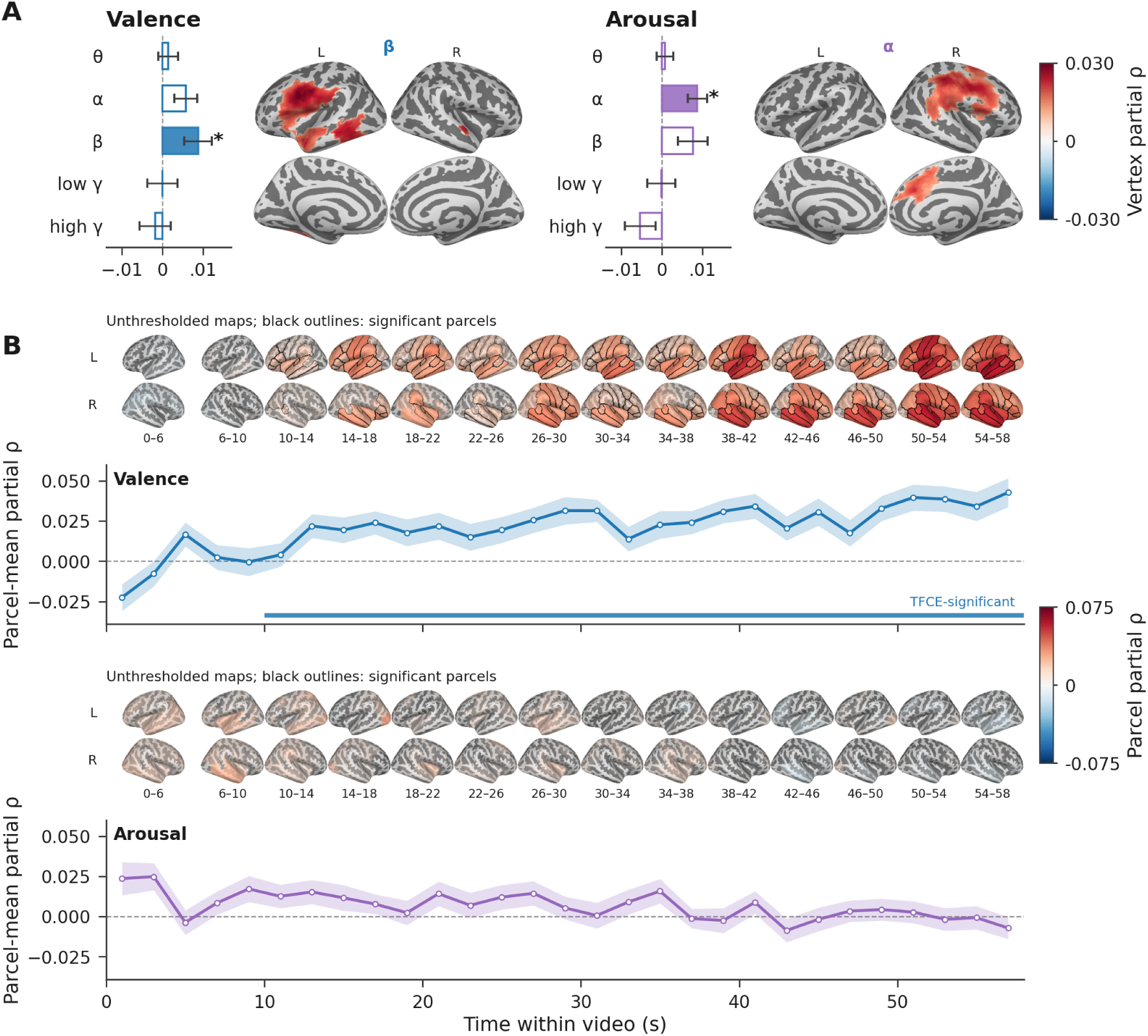
Brain–rating RSA across whole videos and within-video windows. Valence and arousal RDMs were each adjusted for the other affective dimension using partial Spearman correlations. **(A)** Whole-video source spectral RSA in the theta (4–8 Hz), alpha (8–13 Hz), beta (13–30 Hz), low gamma (30–45 Hz), and high gamma (60–90 Hz) bands. Bars show cortex-mean partial ρ ± between-participant SEM (*n* = 92). Filled bars and asterisks identify frequency bands containing at least one vertex surviving spatial TFCE correction; they do not indicate significance tests of the cortex means. Cortical maps show partial ρ at significant vertices in the left (L) and right (R) hemispheres, with lateral views above and medial views below. **(B)** Windowed RSA across 29 consecutive, non-overlapping 2-s windows spanning 0–58 s. Curves show mean partial ρ across cortical parcels, with ribbons indicating between-participant SEM ( *n* = 67–92 across windows). Coloured strips mark windows containing at least one parcel surviving joint space–time TFCE correction. Brain maps show unthresholded partial ρ averaged over the labelled intervals, with left and right lateral views above and below, respectively. Each map summarizes two adjacent windows, except the first, which summarizes three (0–6 s). Black outlines identify parcels significant in at least one constituent window. Both analyses used two-sided participant sign-flip tests with 2,048 permutations and corrected *p* < .05. Correction was applied separately within each model-by-frequency-band map in A and jointly across parcels and windows within each model in B. Stimulus-adjusted results are shown in Supplementary Figure S2.

In the windowed analysis, valence associations survived joint space–time TFCE correction in every 2-s window from 10 to 58 s (minimum TFCE-corrected *p* < .001; Figure 4B). Mean associations over this interval were largest in bilateral superior and middle temporal cortex, while left insular and supramarginal parcels showed significant associations across most windows. Later windows showed broader bilateral involvement. No arousal association survived correction in the windowed analysis (minimum TFCE-corrected *p* = .257). The main whole-video and windowed findings were retained after adjustment for the four stimulus-feature RDMs, with a small additional whole-video arousal–beta association surviving correction (Supplementary Figure S2).

### Physiological representations of valence and arousal

Whole-video physiological dissimilarities were associated with valence in zygomaticus EMG (mean partial ρ = .141, *q* = .001) and corrugator EMG (ρ = .081, *q* = .002), after adjusting for arousal (Figure 5A). After adjusting for valence, arousal was associated with mean positive phasic EDA (ρ = .046, *q* = .039). No other physiological association survived FDR correction.

**Figure 5.**
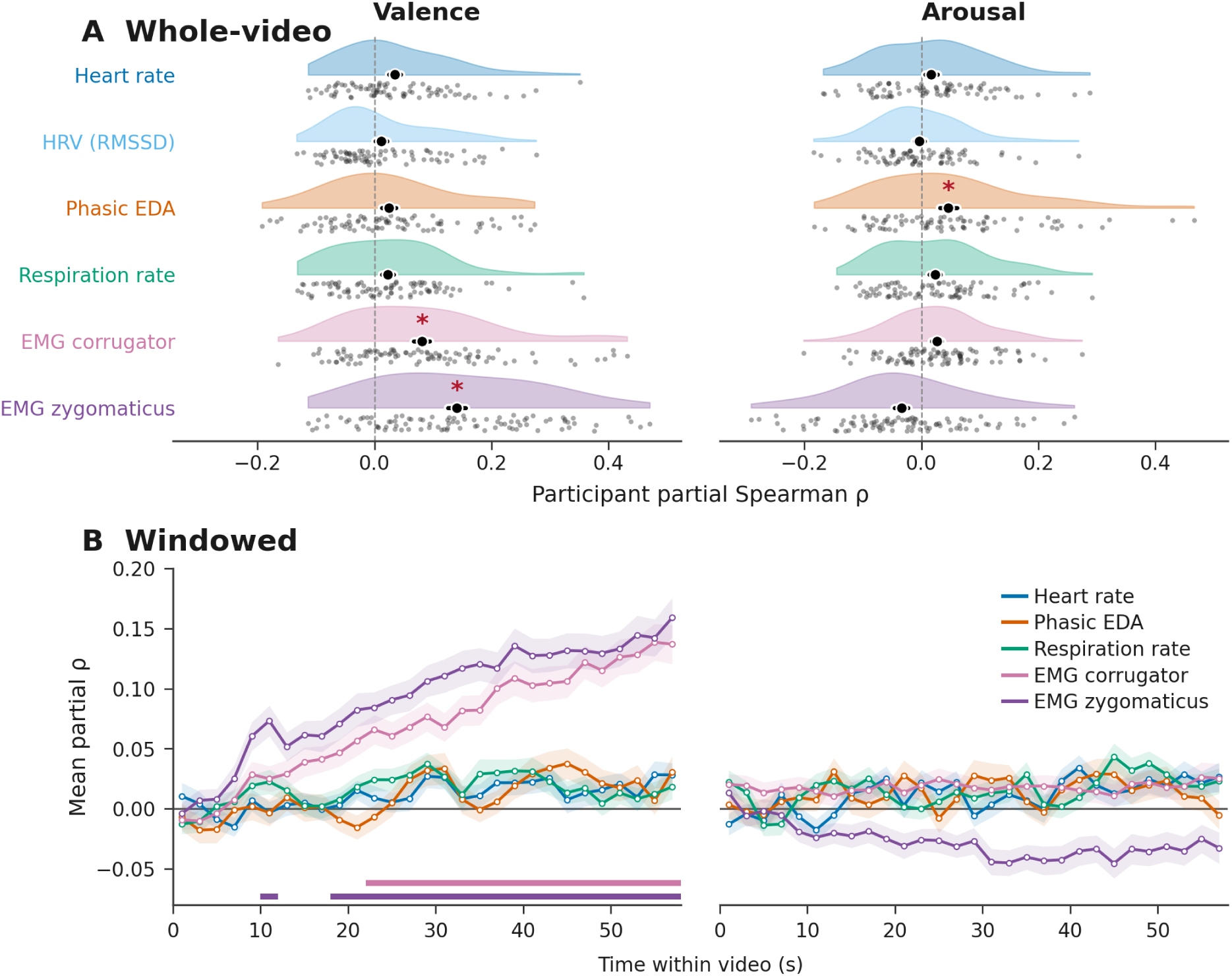
Physiological representations of valence and arousal. Partial Spearman correlations quantify correspondence between physiological and rating RDMs, controlling for the other affective dimension. **(A)** Whole-video associations. Half-violins show participant distributions, grey dots show individual participants, and black markers indicate means ± SEM. Red asterisks identify associations surviving Benjamini–Hochberg FDR correction across the 12 signal-by-rating tests, using 10,000 shared video-label permutations (*q* < .05). **(B)** Windowed associations across 29 consecutive 2-s windows spanning 0–58 s. Lines show participant means and ribbons indicate SEM for all signals. Coloured bars mark windows surviving TFCE correction across time and the five physiological signals within each affective dimension (10,000 shared video-label permutations; two-sided corrected *p* < .05). HRV was evaluated only in the whole-video analysis. Sample sizes were 79 for heart rate and HRV, 80 for EDA, 86 for respiration rate, 92 for corrugator EMG, and 91 for zygomaticus EMG. EDA represents mean positive phasic EDA. Unadjusted associations are shown in Supplementary Figure S4.

Windowed RSA revealed sustained associations between facial-muscle activity and valence after adjusting for arousal (Figure 5B). Associations survived TFCE correction at 10–12 and 18–58 s for zygomaticus EMG (minimum corrected *p* < .001) and at 22–58 s for corrugator EMG (minimum corrected *p* < .001). No autonomic signal showed a significant windowed association with valence, and no physiological signal showed a significant windowed association with arousal.

Within-participant correlations among physiological RDMs were small, with the largest association between heart rate and HRV (mean ρ = .18; Supplementary Figure S3A). Correlations between group-average RDMs were larger, reaching ρ = .65 for heart rate and EDA; their relationships are visualized by MDS (Supplementary Figure S3B,C). Median participant-level VIFs ranged from 1.06 to 1.10, compared with 1.30–2.27 for group-average RDMs (Supplementary Figure S3D). Analyses without adjustment for the other affective dimension retained the whole-video facial-EMG–valence and EDA–arousal associations, with additional associations for heart rate with valence, respiration rate with both dimensions, and corrugator EMG with arousal (Supplementary Figure S4A). Windowed unadjusted analyses likewise showed facial-EMG–valence associations, with no significant arousal associations (Supplementary Figure S4B).

### Temporal correspondence between brain and body

Dynamic brain–body RSA tested whether patterns of differences across videos corresponded between neural and physiological activity sampled at different relative times (Figure 6A). Correspondence exceeded the temporally decoupled reference, obtained by independently sampling physiological times within each video, for corrugator EMG, zygomaticus EMG, heart rate, and phasic EDA, but not respiration rate (Figure 6B–D; minimum corrected *p* = .002 in each significant signal). To summarize the spatial pattern, we ranked parcels by their largest observed-minus-reference correlation at a significant lag and examined the five strongest parcels for each signal. Both facial-muscle signals showed prominent correspondence in insular, anterior frontal, and medial temporal regions. Heart-rate correspondence was strongest in frontal polar and orbital frontal regions and the left insula. EDA correspondence involved left entorhinal and parahippocampal cortex, left insula and frontal pole, and right rostral middle frontal cortex (Figure 6B,C).

**Figure 6.**
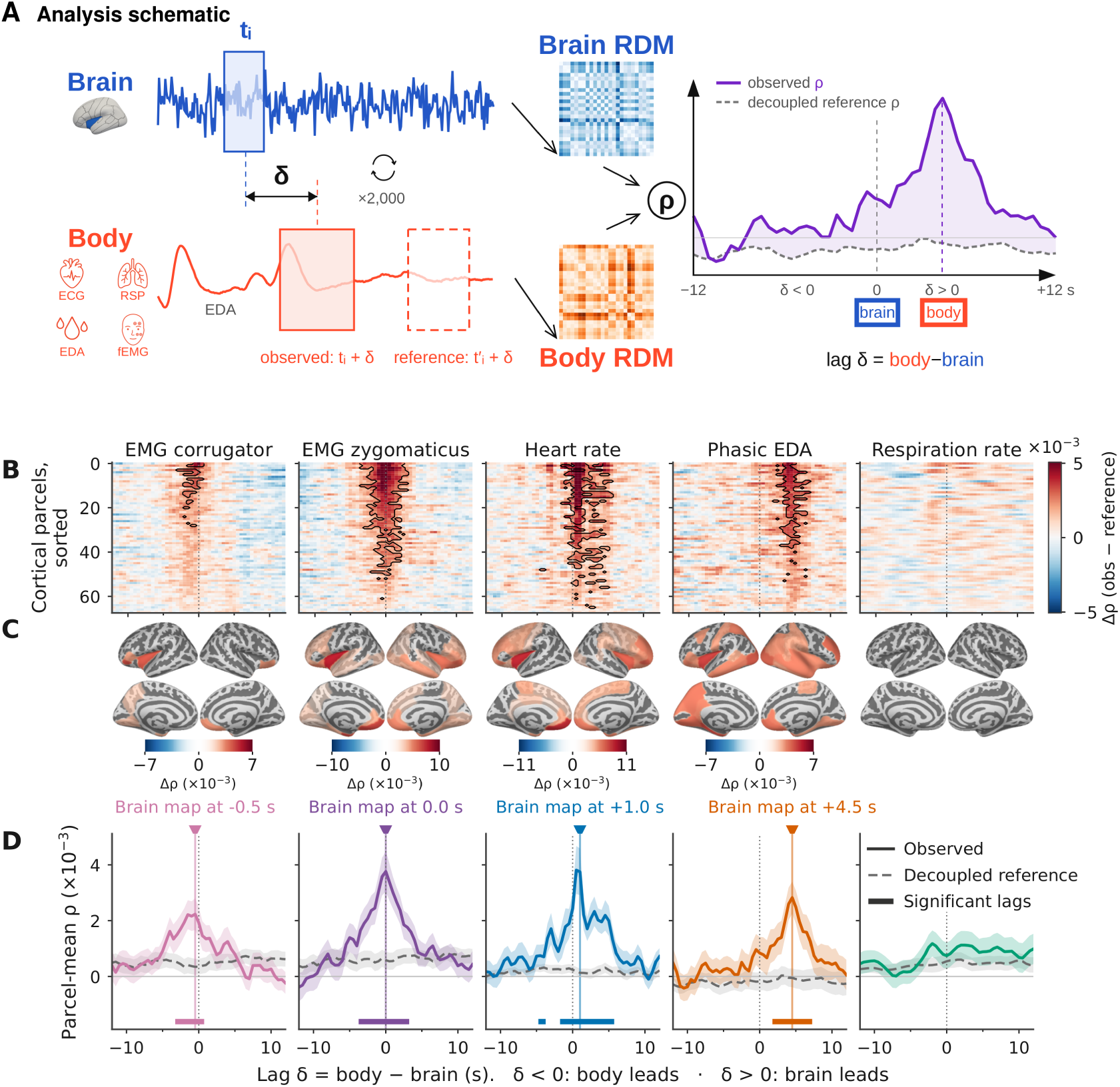
Temporal correspondence between brain and body. **(A)** Analysis schematic. Activity sampled from each of 25 videos forms neural and physiological RDMs, which are compared using Spearman correlation. Physiological sampling times are shifted relative to neural times to examine different lags. Independently sampled physiological times within the same videos provide the temporally decoupled reference. Estimates are averaged over 2,000 repetitions per lag. Example waveforms and RDMs illustrate the procedure; sampling boxes are not drawn to a common duration scale. RSP, respiration; fEMG, facial electromyography. **(B)** Observed minus reference correspondence (Δρ) across 68 cortical parcels and lags. Parcels are sorted independently for each signal by their maximum absolute Δρ. Black contours identify significant parcel–lag combinations. Lag values are labelled in panel D. **(C)** Significant cortical effects at the map times marked above panel D, selected at the maximum parcel-mean Δρ for each signal. Lateral views are shown above medial views. Each signal has its own colour scale, expressed as Δρ × 10⁻³. **(D)** Parcel-mean observed and reference correlations, with ribbons indicating between-participant SEM. Coloured bars mark lags containing significant effects. Triangles identify the times selected for the cortical maps, rather than the weighted-average lags reported in the text. “Body leads” denotes earlier physiological sampling and “Brain leads” denotes later physiological sampling; these labels indicate temporal ordering, not causal direction. Significance was assessed using two-sided TFCE permutation tests with 2,048 permutations, corrected across parcels and lags within each signal and across five signals using Bonferroni correction (corrected *p* < .05). Sample sizes were 91 for corrugator, 90 for zygomaticus, 78 for heart rate, 79 for EDA, and 85 for respiration rate.

The temporal profiles differed across signals. Summarized by the weighted-average lag of significant effects, corrugator correspondence occurred at −1.0 s (95% CI [−1.1, −0.9] s), zygomaticus at 0.0 s (95% CI [−0.1, 0.2] s), heart rate at +1.7 s (95% CI [1.5, 1.9] s), and EDA at +4.5 s (95% CI [4.4, 4.6] s). Negative lags indicate that physiological activity occurred earlier than neural activity; positive lags indicate that it occurred later. These values summarize temporal correspondence rather than causal transmission delays.

## Discussion

Using a common representational framework, we examined how subjective experience, neural activity and peripheral physiology relate during emotional video viewing. Across whole videos, valence-related similarity was associated with beta-band neural power patterns and facial-muscle responses, whereas arousal-related similarity was associated with alpha-band power patterns and phasic electrodermal activity. Comparisons within successive 2-s windows showed that videos rated similarly in valence also elicited similar neural patterns and facial-muscle responses across much of the viewing period, whereas no significant windowed associations were detected for arousal. Dynamic RSA further revealed temporal correspondence between brain and body response patterns, with facial-muscle correspondence centred near zero lag and cardiac and electrodermal correspondence centred at later physiological times. Together, these findings link subjective affect to complementary neural and bodily response patterns and show that brain–body correspondence follows different temporal profiles across physiological systems.

The neural findings support a representational account in which similarities between emotional experiences are reflected in similarities between cortical response patterns. Previous fMRI studies identified valence-related patterns that generalized across sensory modalities (Chikazoe et al., 2014) and correspondence between neural and subjective similarities among emotional states (Saarimäki et al., 2016). Our findings extend this account to electrophysiological activity during sustained viewing, with valence-related similarity expressed in beta-band patterns, predominantly in left sensorimotor and frontotemporal cortex. This distribution is compatible with MEG evidence linking dynamic facial-expression recognition to beta activity in superior temporal, sensorimotor and frontolimbic regions (Jabbi et al., 2015). These observations suggest that perceptual and sensorimotor processes may contribute to the evaluation of emotional scenes. Preservation of the associations after adjustment for measured visual and semantic similarities further supports their relationship with affective evaluations, although other unquantified aspects of scene content may also contribute.

The alpha–arousal association complements evidence that emotional intensity modulates low-frequency neural activity. Highly arousing pleasant and unpleasant pictures induce alpha and lower-beta suppression (Schubring & Schupp, 2021), while continuous arousal during movie viewing covaries with low-frequency EEG power (Camenzind et al., 2026). Our result links arousal to the spatial organization of alpha power, complementing these amplitude-based findings. Its right frontoparietal and sensorimotor distribution suggests that attentional and sensorimotor engagement may contribute to this correspondence. Sensorimotor alpha is itself sensitive to affective context, including observing pain in others (Whitmarsh et al., 2011). These convergences provide functional context for the spectral findings without implying exclusive mappings of beta to valence or alpha to arousal.

The windowed results suggest that correspondence with the overall valence evaluation extends across much of the viewing period. Significant associations were detected in successive windows from 10 to 58 s, with the cortex-averaged association largest near the end. One possible explanation is that viewers initially accumulate contextual information to establish an overall positive or negative interpretation, which remains reflected in neural responses as the scene unfolds. Naturalistic film studies show that responses in association cortex depend on information integrated over several seconds, consistent with this context-building account (Hasson et al., 2008). The first significant window nevertheless does not identify a minimum time required to recognize valence. Stronger correspondence near the end may also reflect how experiences are summarized retrospectively: experimental studies show that final moments can receive disproportionate weight in overall emotional evaluations (Fredrickson & Kahneman, 1993; Li & Lapate, 2024). The observed time course could therefore reflect both accumulating context and the weighting of recent experience in the subsequent rating.

Arousal-related correspondence may depend more strongly on the timing of individual events. If arousing moments occur at different points across clips, comparing the same elapsed 2-s interval in every video with a single overall intensity rating could miss relevant responses occurring elsewhere in each clip. Studies using time-varying arousal ratings have detected neural associations during movie viewing, including associations with dynamic functional connectivity that generalize across films (Ke et al., 2025). Our windowed arousal null could therefore reflect a mismatch between momentary responses and retrospective evaluation. Together, the findings suggest that valence-related correspondence is relatively sustained during viewing, whereas arousal-related patterns may be more transient and tied to particular events. Continuous ratings and event-aligned analyses would allow this distinction to be tested directly.

The facial-muscle findings connect retrospective valence judgments with expressive responses across much of video viewing. Pleasant stimuli typically increase zygomaticus activity and reduce corrugator activity, although the two muscles have different response functions (Lang et al., 1993; Larsen et al., 2003). Because RSA compares differences between videos, these opposite amplitude relationships can both produce positive correspondence with valence. Film studies also link changes in subjective valence to facial EMG (Golland et al., 2018; Sato et al., 2020), with closer correspondence during emotional than neutral viewing (Constantinou et al., 2023). Our results extend this work by relating facial responses during viewing to the subsequent overall evaluation. Correspondence with retrospective valence was sustained across successive windows and generally strengthened as the videos progressed, reaching its largest values near the end (Figure 5B). Thus, facial responses increasingly distinguished the clips in a way that matched participants’ eventual valence judgments. Alongside the neural findings, this pattern links the final evaluation to responses expressed across much of the emotional episode, with particularly close correspondence near its conclusion.

The modest whole-video association between mean positive phasic EDA and arousal is consistent with evidence linking skin-conductance response magnitude or frequency to subjective arousal (Bradley et al., 2001; Lang et al., 1993; Xia et al., 2017). Heart rate, heart-rate variability and respiration rate showed no corrected associations after accounting for the other affective dimension. These analyses ask whether differences between videos in each physiological summary match differences in their overall valence or arousal ratings. The selected features capture only part of autonomic responding: for example, arousal during film viewing can alter expiratory timing, airflow and ventilation, which respiration rate alone does not fully describe (Gomez et al., 2005). The modest EDA association and limited cardiac and respiratory correspondence are therefore compatible with partial overlap between subjective and autonomic responses. Such overlap was demonstrated by Cuve et al. (2023), whose analysis identified both autonomic variation shared with subjective valence and arousal and additional variation not represented in those ratings.

Dynamic RSA revealed that momentary differences between videos in cortical activity corresponded to differences in facial-muscle, cardiac and electrodermal responses. This correspondence exceeded the temporally decoupled reference, showing that the relative timing of neural and physiological fluctuations mattered. Building on dynamic RSA developed for naturalistic vision (de Vries & Wurm, 2023), our analysis compares cortical activity with multiple physiological signals within a common representational similarity framework. Heart-rate correspondence was detected despite the absence of corrected associations with retrospective valence or arousal ratings. This finding illustrates the complementary perspective provided by examining ongoing brain–body relationships alongside associations with an episode’s overall evaluation.

The spatial findings suggest partially shared cortical involvement across physiological systems. The left insula was among the five strongest parcels for both facial muscles, heart rate and EDA, with additional prominent associations in frontal and medial temporal cortex. This convergence has several precedents. Electrical stimulation of the human insula can produce cardiac acceleration or deceleration (Chouchou et al., 2019), and intracranial recordings during movie viewing link changing insula–cingulate interactions to heart-rate variability (Sonkusare et al., 2023). Simultaneous facial EMG and fMRI have also associated facial responses with inferior frontal activity, with corrugator responses additionally related to insular and hippocampal activity (Likowski et al., 2012). Our findings extend this convergence across expressive and autonomic responses recorded during the same emotional episodes. The overlapping anatomy is consistent with distributed systems that monitor and regulate bodily states (Kleckner et al., 2017; Zhang et al., 2025), although the spatially extensive effects do not identify specialized generators for individual signals, and MEG source estimates have limited spatial specificity for deep regions such as the insula and medial temporal cortex.

Facial correspondence occurred relatively close to the neural fluctuations, with weighted-average lags of approximately −1 s for corrugator and zero for zygomaticus. Facial responses can develop rapidly: EMG differentiates reactions to happy and angry expressions within approximately 300– 400 ms of stimulus presentation (Dimberg & Thunberg, 1998). Facial activity can also influence central processing. Experimentally reducing frown-muscle feedback attenuated amygdala responses during deliberate imitation of angry expressions (Hennenlotter et al., 2009). These observations support considering facial expression and neural processing as interacting components of an unfolding emotional response. Our temporal profiles are compatible with this reciprocal account, but the earlier corrugator correspondence alone does not establish a feedback mechanism. Residual facial-muscle contamination of MEG may also contribute to the observed association.

Cardiac correspondence was weighted toward later physiological times, with an average lag of approximately +1.7 s, although significant effects extended across zero. Part of this lag is expected from the measure itself: heart rate is derived from completed interbeat intervals, and cardiac responses to an event develop over several beats. This broad profile should be understood in the context of reciprocal brain–heart interactions. Model-based analyses of EEG and heart-rate variability have identified ascending cardiac influences at the onset of emotional arousal, followed by bidirectional interactions (Candia-Rivera et al., 2022). Our analysis addresses a different aspect of this relationship: the relative timing of similarities across videos in heart rate and cortical activity, summarized across viewing. The positive average lag therefore indicates that corresponding heart-rate patterns occurred later than neural patterns, but does not establish a causal direction.

EDA showed a later profile, with significant correspondence at positive lags and an average lag of approximately +4.5 s. The slow development of skin-conductance responses provides a plausible physiological explanation. Empirical response models show that skin conductance unfolds over several seconds, with its timing shaped by the peripheral response and recording location (Bach et al., 2010). Earlier MEG work also linked emotional enhancement of an early occipitotemporal response to the magnitude of subsequent skin-conductance responses across participants (D’Hondt et al., 2010). Our findings extend this general relationship to ongoing phasic fluctuations during video viewing. The observed lag plausibly combines the timing of cortical–autonomic relationships with the slower expression of electrodermal activity, and should not be equated with a neural transmission delay.

The absence of corrected correspondence for respiration rate also deserves a feature-specific interpretation. MEG research has demonstrated that respiratory phase modulates cortical oscillatory activity (Kluger & Gross, 2021); our analysis instead asked whether differences in breathing rate matched differences in cortical activity. Together, the dynamic findings reveal overlapping cortical involvement alongside distinct temporal profiles across bodily systems. These relationships may reflect both physiological interactions and coordinated responses to unfolding film events. During informative video viewing, Madsen and Parra (2022) found coupling between neural, cardiac and ocular activity within individuals, alongside synchrony across viewers that varied with attention. Attentional engagement may therefore contribute to the brain–body correspondence observed during emotional viewing. Because affective ratings were not included in dynamic RSA, the findings establish temporally structured brain–body correspondence during emotional viewing without specifying whether that correspondence represents valence, arousal or other aspects of the experience.

Several limitations should be considered when interpreting the present findings. First, all participants viewed the 25 clips in the same sequence, so the design cannot separate a clip’s content from its position in the session or from the influence of preceding clips. Rating dissimilarities showed little relationship with the temporal separation between clips (group-average ρ = −.07 for valence and −.02 for arousal; Supplementary Figure S1B); a gradual change in neural or bodily state across the session would therefore not be expected to resemble the rating structure. Replication with a broader stimulus set and presentation orders varied across participants would be needed to distinguish these contextual influences from responses to the clips’ content and to assess generalizability (Schütt et al., 2023). Second, post-video ratings captured overall valence and arousal, providing a summary of each emotional episode.

Continuous measures of experienced affect could complement these overall evaluations by clarifying how neural and bodily response patterns relate to changes in subjective experience during viewing. Finally, the spectral analyses used band-limited power, which can reflect both oscillatory activity and changes in the aperiodic background (Donoghue et al., 2020). Residual muscle activity may also contribute to MEG estimates, particularly at higher frequencies, despite the artifact-reduction procedures. Further separation of oscillatory and aperiodic components, together with assessment of residual muscle contributions, would help refine the physiological interpretation of the spectral associations.

In conclusion, this study brings together two complementary perspectives on emotional experience: how an episode is evaluated as a whole and how neural and bodily responses relate during its unfolding. Examining both within a common representational similarity framework reveals shared organization alongside distinct temporal relationships across emotional components. These findings support a multicomponent view of emotion in which subjective evaluations, cortical activity and peripheral physiology provide partly overlapping perspectives on the same experience. Understanding their coordination therefore involves considering both which aspects of an experience they reflect and when their responses correspond. This approach provides a foundation for investigating how ongoing brain–body dynamics relate to the emotional meaning people assign to their experiences.

## Data and code availability statement

The MEG and physiological data are not publicly available due to legal restrictions. The analysis code is available in the [GitHub repository](https://github.com/weiyongxu/video-task-analysis).

## Author contributions

Weiyong Xu: Methodology; Data curation; Formal analysis; Software; Validation; Visualization; Writing – original draft; Writing – review & editing.

Pinja Ronkainen: Conceptualization; Methodology; Investigation; Data curation; Validation; Writing – review & editing.

Tiina Parviainen: Conceptualization; Methodology; Resources; Funding acquisition; Project administration; Supervision; Writing – review & editing.

## Acknowledgments

We thank Kristina Pultsina for the help with physiological signal processing.

