## Supplemental Data 1 for "Affective representations and brain–body dynamics during emotional video viewing"

#### Supplementary methods: post-video ratings

The questions were presented in Finnish after each video. Original Finnish wording is followed below by its English translation. For the five discrete-emotion questions, scale endpoints were 0 = ei lainkaan (not at all) and 7 = hyvin paljon (very much).

- **Familiarity.** "Oletko nähnyt videon aiemmin?" — "Have you seen the video before?" Responses: Kyllä / Ei (yes / no).
- **Categorical valence.** "Oliko video mielestäsi kokonaisuudessaan negatiivinen, positiivinen vai neutraali?" — "In your opinion, was the video overall negative, positive or neutral?" Responses: Negatiivinen / Positiivinen / Neutraali (negative / positive / neutral).
- **Valence.** "Asteikolla 0-7, miltä video sinusta tuntui?" — "On a scale of 0–7, how did the video make you feel?" Scale endpoints: 0 = hyvin epämiellyttävä (very unpleasant) and 7 = hyvin miellyttävä (very pleasant).
- **Intensity (arousal).** "Kuinka intensiivinen video oli?" — "How intense was the video?" Scale endpoints: 0 = ei lainkaan (not at all) and 7 = hyvin intensiivinen (very intense).
- **Happiness.** "Asteikolla 0-7, kuinka vahvasti koit videota katsoessasi: ILOA?" — "On a scale of 0–7, how strongly did you experience happiness while watching the video?"
- **Contentment.** "Asteikolla 0-7, kuinka vahvasti koit videota katsoessasi: TYYTYVÄISYYTTÄ?" — "On a scale of 0–7, how strongly did you experience contentment while watching the video?"
- **Anxiety.** "Asteikolla 0-7, kuinka vahvasti koit videota katsoessasi: AHDISTUSTA?" — "On a scale of 0–7, how strongly did you experience anxiety while watching the video?"
- **Fear.** "Asteikolla 0-7, kuinka vahvasti koit videota katsoessasi: PELKOA?" — "On a scale of 0–7, how strongly did you experience fear while watching the video?"
- **Sadness.** "Asteikolla 0-7, kuinka vahvasti koit videota katsoessasi: SURUA?" — "On a scale of 0–7, how strongly did you experience sadness while watching the video?"

#### Supplementary methods: stimulus selection criteria

The final validation survey included 16 respondents. Selection criteria were based on intensity, valence, and target-emotion ratings averaged across the videos within each intended emotion category. For valence, the criteria were mean ratings below 3 for negative videos (anxiety/fear and sadness), between 3 and 4 for neutral videos, and above 4 for positive videos (happiness and contentment). Intensity ratings were mainly used to check that happiness videos were more intense than contentment videos and that anxiety/fear videos were more intense than sadness videos; both distinctions were supported by the validation ratings. Neutral videos were also required to have a mean intensity rating below 2. For the target emotion, the criterion was the highest mean rating among the assessed emotions—for example, the highest rating for anxiety in anxiety/fear videos. Mean contentment ratings were the same for contentment and happiness videos. Similar overlap between happiness and contentment ratings has been reported for a film intended to elicit contentment (Gross & Levenson, 1995). Both categories were retained because their different intensity levels were considered sufficient to distinguish the intended categories.

### Supplementary figures

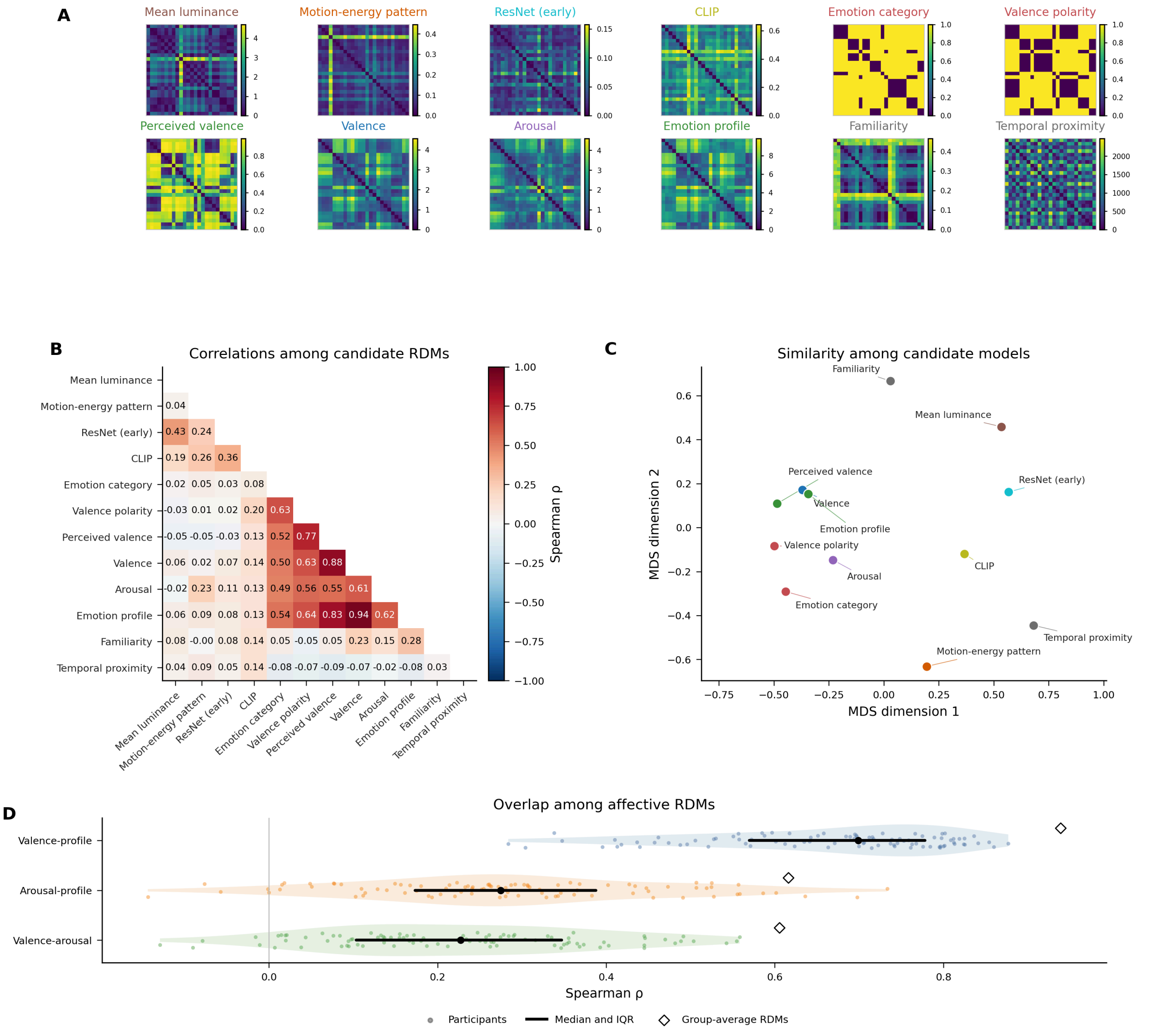


**Supplementary Figure S1. Candidate stimulus and rating RDMs. (A)** Representational dissimilarity matrices for 25 videos, describing stimulus features, categorical models, subjective ratings, familiarity, and temporal proximity. The categorical models encode the intended emotion category (anxiety/fear, sadness, neutral, contentment, or happiness), intended valence polarity (negative, neutral, or positive), and each participant's perceived-valence judgment (negative, neutral, or positive). For these models, dissimilarity is 0 for videos in the same category and 1 for videos in different categories. Participant-specific RDMs were averaged across 92 participants. Rows and columns follow the same video order across matrices; colour scales are specific to each model. **(B)** Spearman correlations across the 300 unique video pairs in the group-average RDMs, with self-correlations omitted. **(C)** Two-dimensional metric multidimensional scaling (MDS) of model dissimilarities, defined as $1 - \rho$ from panel B. Distances approximate model dissimilarities; axis orientation and the origin are arbitrary (normalized Stress-1 $= 0.194$). Colours for the six models used in stimulus-adjusted RSA match Figure 3. Emotion category and valence polarity are shown in red, perceived valence and emotion profile in green, and familiarity and temporal proximity in grey. **(D)** Participant-level Spearman correlations among valence, arousal, and emotion-profile RDMs. Dots represent participants ($n = 92$), pale shading shows their distributions, and black points and bars indicate medians and interquartile ranges. Open diamonds show correlations between group-average RDMs rather than averages of participant-level correlations. "Profile" denotes the RDM constructed from the five discrete-emotion ratings.

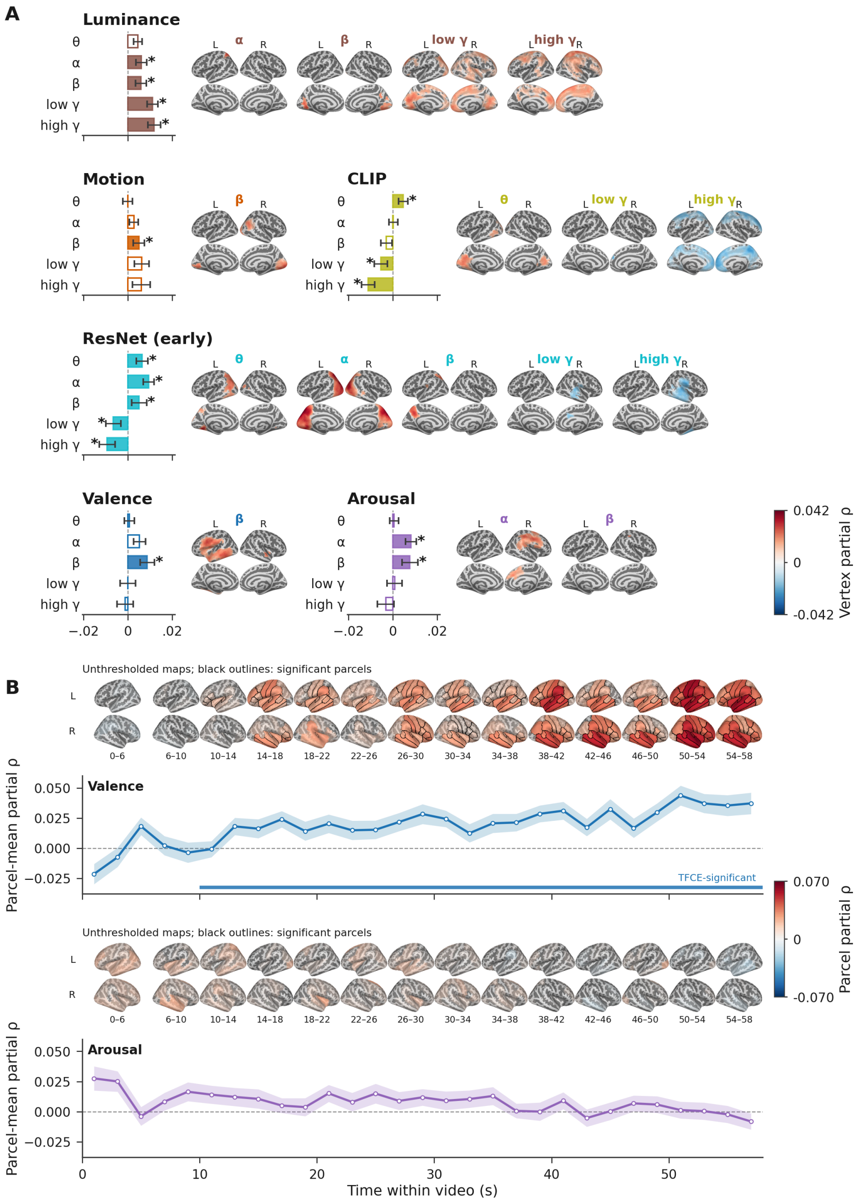


**Supplementary Figure S2. Brain–rating RSA adjusted for measured stimulus features. (A)** Whole-video source spectral RSA for six models: mean luminance, motion-energy patterns, early visual-network features (ResNet), high-level semantic features (CLIP), valence, and arousal. Each model was adjusted for the other five using partial Spearman correlations. Bars show cortex-mean partial ρ ± between-participant SEM ($n = 92$). Filled bars and asterisks identify frequency bands containing at least one vertex surviving spatial TFCE correction; they do not indicate significance tests of the cortex means. Maps show partial ρ at significant vertices in the left (L) and right (R) hemispheres, with lateral views above and medial views below. Bands are theta (4–8 Hz), alpha (8–13 Hz), beta (13–30 Hz), low gamma (30–45 Hz), and high gamma (60–90 Hz). **(B)** Windowed valence and arousal RSA with the same adjustments across 29 consecutive 2-s windows spanning 0–58 s. Curves show parcel-mean partial ρ, with ribbons indicating between-participant SEM ($n = 67$–$92$ across windows). Coloured strips mark windows containing at least one parcel surviving joint space–time TFCE correction. Maps show unthresholded partial ρ averaged over the labelled intervals, with left and right lateral views above and below. Each map summarizes two adjacent windows, except the first (0–6 s), which summarizes three. Black outlines identify parcels significant in at least one constituent window. Both analyses used two-sided participant sign-flip tests with 2,048 permutations and corrected $p < .05$. Correction was applied separately within each model-by-frequency-band map in A and jointly across parcels and windows within each model in B. Colour scales differ between panels.

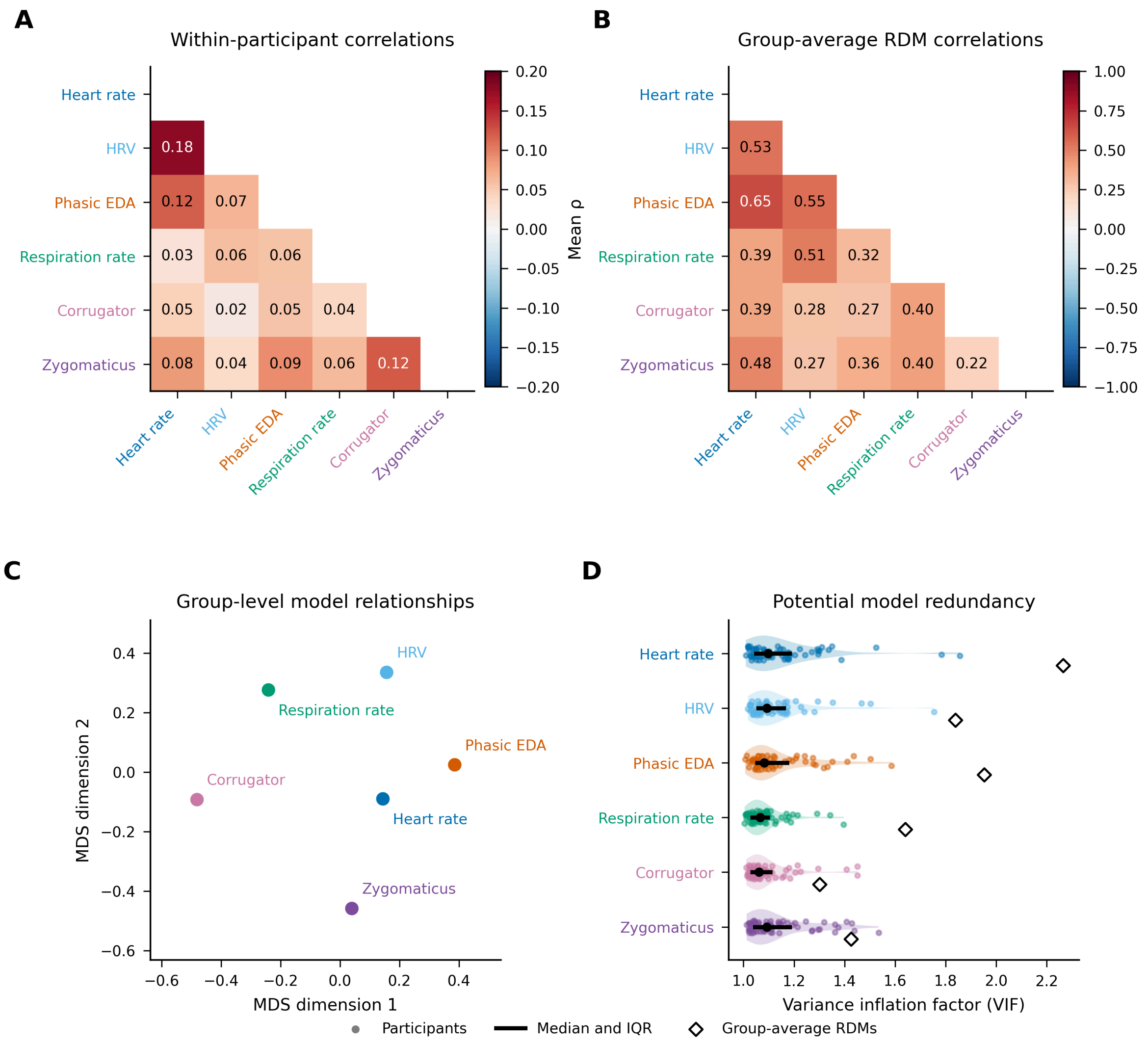


**Supplementary Figure S3. Relationships among physiological RDMs.** Six physiological features correspond to those used in the whole-video body–rating analyses: heart rate, HRV (RMSSD), mean positive phasic EDA, respiration rate, and corrugator and zygomaticus EMG envelopes. **(A)** Within-participant Spearman correlations between RDMs, averaged in Fisher-$z$ space and back-transformed. Each pair uses participants with both signals available and their shared finite video pairs. **(B)** Spearman correlations between group-average RDMs, calculated after standardizing each participant's 300 unique video-pair dissimilarities within each signal and averaging across participants. Self-correlations are omitted in A and B; colour scales differ. **(C)** Two-dimensional metric multidimensional scaling (MDS) of $1 - \rho$ from B. Nearby signals have more similar group-average RDMs; axis orientation is arbitrary (normalized Stress-1 $= 0.168$). **(D)** Variance inflation factors (VIFs) calculated from ranked RDM vectors for the 63 participants with all six signals available. Dots show participants, violins show distributions, black markers and bars show medians and interquartile ranges, and open diamonds show VIFs calculated from group-average RDMs. VIFs describe potential redundancy in a joint physiological model; the reported body–rating analyses tested one physiological RDM at a time. Signal-specific sample sizes for group-average RDMs were 79 for heart rate and HRV, 80 for EDA, 86 for respiration rate, 92 for corrugator EMG, and 91 for zygomaticus EMG. Panels A–D are descriptive.

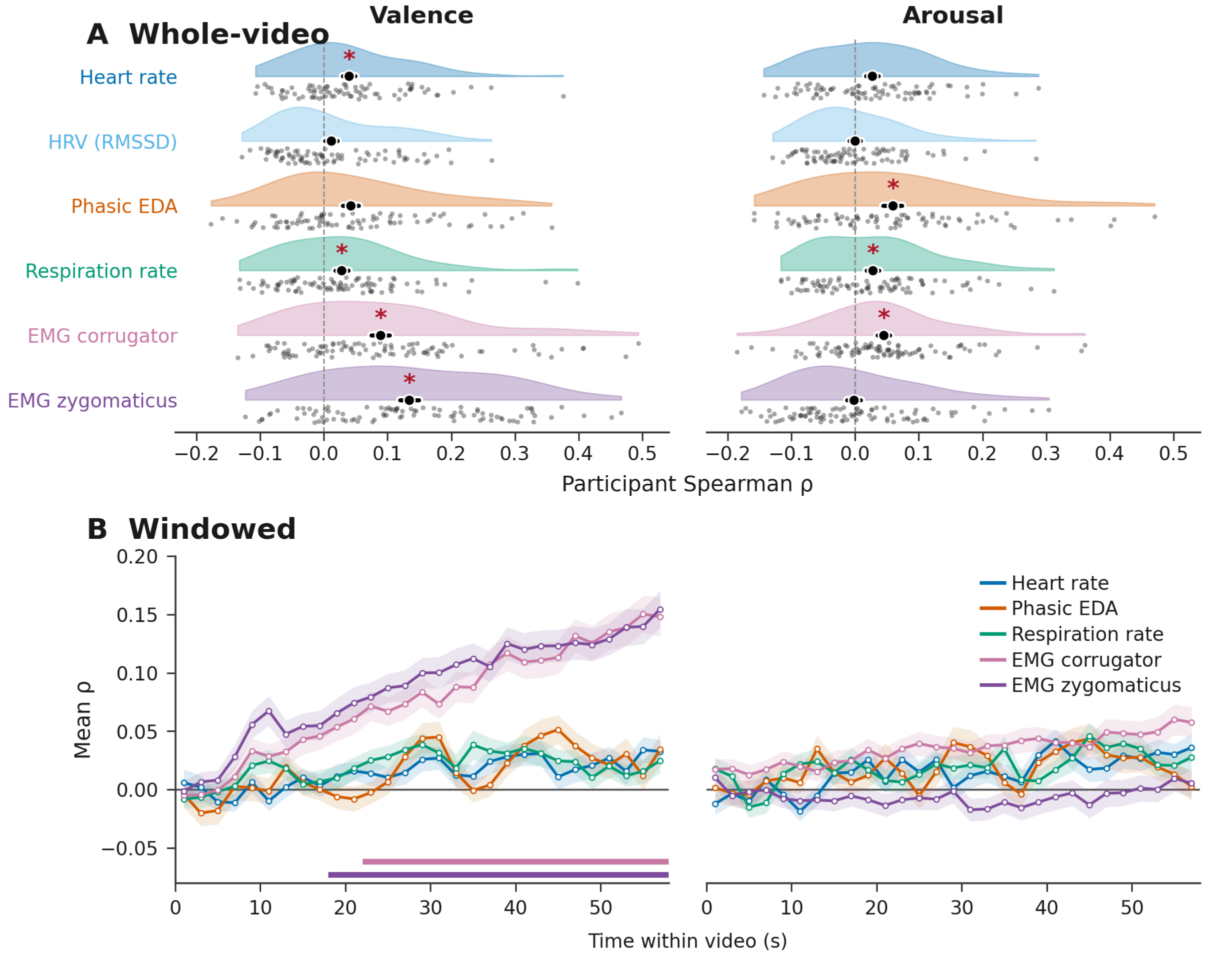


**Supplementary Figure S4. Physiological associations without adjustment for the other affective dimension.** Spearman correlations quantify correspondence between physiological and valence or arousal RDMs without controlling for the other rating dimension. **(A)** Whole-video associations. Half-violins show participant distributions, grey dots show individual participants, and black markers indicate means ± SEM. Red asterisks identify associations surviving Benjamini–Hochberg FDR correction across the 12 signal-by-rating tests, using 10,000 shared video-label permutations ($q < .05$). **(B)** Windowed associations across 29 consecutive 2-s windows spanning 0–58 s. Lines show participant means and ribbons indicate SEM for all signals. Coloured bars mark windows surviving TFCE correction across time and the five physiological signals within each affective dimension (10,000 shared video-label permutations; two-sided corrected $p < .05$). HRV was evaluated only in the whole-video analysis. Sample sizes were 79 for heart rate and HRV, 80 for EDA, 86 for respiration rate, 92 for corrugator EMG, and 91 for zygomaticus EMG. EDA represents mean positive phasic EDA. Figure 5 shows the corresponding analyses adjusted for the other affective dimension.
